# The evolution of no-cost resistance at sub-MIC concentrations of streptomycin in *Streptomyces coelicolor*

**DOI:** 10.1101/062414

**Authors:** Sanne Westhoff, Tim M. van Leeuwe, Omar I. Qachach, Zheren Zhang, Gilles P. van Wezel, Daniel E. Rozen

**Author notes:** These authors contributed equally to this work (Sanne Westhoff, Tim M. van Leeuwe). Corresponding author: Daniel E. Rozene Institute of Biology Leiden Sylviusweg2333 BE Leiden.

## Abstract

At the high concentrations used in medicine, antibiotics exert strong selection on bacterial populations for the evolution of resistance. However, these lethal concentrations may not be representative of the concentrations bacteria face in soil, a recognition that has lead to questions of the role of antibiotics in soil environments as well as the dynamics of resistance evolution during sub-lethal challenge. Here we examine the evolution of resistance to sub-MIC concentrations of streptomycin in the filamentous soil bacterium *Streptomyces coelicolor*. First, we show that spontaneous resistance to streptomycin causes an average fitness deficit of ~21% in the absence of drugs; however, these costs are eliminated at concentrations as low as 1/10 the MIC of susceptible strains. Using experimental evolution, we next show that resistance readily evolves at these non-lethal doses. More important, *S. coelicolor* resistance that evolves at sub-MIC streptomycin is cost-free. Whole-genome analyses reveal that sub-MIC evolved clones fix a distinct set of mutations to those isolated at high drug concentrations. Our results broaden the conditions under which resistance can evolve in nature and suggest that the long-term persistence of these strains is facilitated by the absence of pleiotropic fitness costs. Finally, our data cast doubt on arguments that low-concentration antibiotics in nature are signals, instead supporting models that resistance evolves in response to antibiotics used as weapons.

## Introduction

Because of their lethal effects on target bacteria, antibiotics exert strong natural selection on bacterial populations for the evolution of resistance ^1^. At the high concentrations used in clinical environments, antibiotic resistant clones can rapidly increase in frequency because these strains gain an absolute advantage compared to their susceptible counterparts ^2^. However, it is likely that these high concentrations, above the so-called mutant selection window ^3^, represent an extreme of the drug concentrations bacteria naturally experience ^4^. Drug concentrations within patients can vary markedly through time and across body sites due to difference in drug penetrance, excretion or metabolism ^5,6^. Equally, in the natural environment, where environmental bacteria are exposed to antibiotics from anthropogenic sources as well as endogenous antibiotics produced by bacteria and fungi, bacteria may experience a broad range of drug concentrations ^7,8^. For example, exposure to anthropogenic sources of antibiotics will be greatest near the point of contamination and declines with distance from this source. And although the overall drug concentrations due to endogenous sources are likely low ^9^, gradients in concentrations are anticipated as a function of the distance from these antibiotic-producing microbes. While decades of research have unraveled the dynamics of the evolution of antibiotic resistance at high drug concentrations, scarcely little is understood of the emergence of resistance at the low concentrations that are more reflective of natural values ^10^. What are the dynamics of resistance evolution at low antibiotic concentrations outside of the traditional mutant selective window? And if resistance evolves, is it associated with the same pleiotropic costs borne by clones that evolve resistance after exposure to high drug concentrations? Here we address these questions with a focus on the evolution of streptomycin resistance in the environmental bacterium *Streptomyces coelicolor*. Streptomycetes produce a wide range of natural products, including some 50% of all known antibiotics ^11,12^, and are also well-known environmental reservoirs of antimicrobial resistance ^13^; they are therefore ideal organisms for this study.

Pharmacodynamic models assume that drug-resistant mutants are selected when antibiotic concentrations fall into a specific range known as the mutant selection window ^3,8,14^. This traditional selective window encompasses the antibiotic concentrations between the minimal inhibitory concentration (MIC) of the susceptible strain and the MIC of the resistant strain ^14^. However, while this model correctly identifies the MIC as the threshold where resistant cells persist and susceptible cells die, it fails to account for the fact that below the MIC the two cell types are not otherwise competitively equivalent ^8^. Indeed, susceptible cells can be significantly harmed by non-lethal, sub-MIC, antibiotics and these negative effects on growth can markedly increase the range of drug concentrations where resistant cells are selected ^6^. Of equal importance, the antibiotic concentration where resistance evolves can have crucial implications for the type of resistance that evolves ^8,15^.

While antibiotic resistance that evolves at high concentrations often has a significant cost in terms of bacterial fitness ^1^, recent studies have predicted that this cost will not be evident for resistance that emerges at low drug concentrations ^6,8,16^. The reasons for this can be intuitively explained as follows: while resistant cells above the MIC gain an absolute fitness advantage against susceptible strains, below the MIC, resistant cells and susceptible cells will compete with one another. Accordingly, the success of resistant strains below the MIC will be determined both by their ability to withstand the effects of drug exposure and also their intrinsic competitiveness relative to susceptible cells. Strains with costly resistance may therefore fail to outcompete susceptible strains, while strains with no-cost resistance will thrive. As a consequence of these lower costs, it is furthermore predicted that resistance that evolves at sub-MIC antibiotic concentrations will persist when growing in environments without drugs, whereas strains with costly resistance may be outcompeted ^17^.

Our aims here are to quantify the concentration dependent fitness effects of spontaneous streptomycin resistance in *S. coelicolor*. Streptomycin is an aminoglycoside antibiotic that is produced in the soil by the natural antibiotic producer *Streptomyces griseus* ^18^; although difficult to directly quantify, it is believed that streptomycin concentrations in soil are extremely low, raising questions about the role of this antibiotic in nature for the bacteria that produce it ^18^. It has even been argued that because antibiotics at such low, non-lethal, concentrations are insufficient to select for resistance, these secondary metabolites are better viewed as signals than as weapons ^9, 19,20^. The results of the present work fail to support this perspective. We first show that rates of streptomycin resistance among natural bacterial isolates are relatively high. Next, we show that while resistance that evolves at high concentrations of antibiotics is highly costly, resistance evolving at sub-MIC drug concentrations is cost-free. We discuss the implications of these results for understanding the evolution and persistence of resistant bacterial strains in nature and also for understanding the roles of antibiotics in natural environments.

## Materials and Methods

### Bacterial strains and culturing conditions

Two *Streptomyces coelicolor* strains were used in this study: *S. coelicolor* A3(2) M145 (designated WT) and *S. coelicolor* A3(2) M145 Apra, an isogenic strain carrying an integrated pSET152 plasmid conferring apramycin resistance (designated WT_Apr_). The MIC of streptomycin for both ancestral strains is 2 μg ml^−1^, indicating that there is no cross-resistance between apramycin and streptomycin (methods for MIC determination are outlined below). Strains were routinely grown at 30 °C on Soy Flour Mannitol Agar (SFM) containing 20 g Soy Flour (Biofresh, Belgium), 20 g Mannitol (Merck KGaA, Germany) and 15 g agar (Hispanagar, Spain) per liter (pH 7.2 - 7.4) To generate high-density spore stocks, plates were uniformly spread with 50 μl of spore containing solution. After 3-4 days of growth, spores were harvested with a cotton disc soaked in 3 ml 30% glycerol, and then spores were extracted from the cotton by passing the liquid through an 18g syringe to remove the vegetative mycelium. Resulting spore stocks were titred and stored at −20 °C. Growth rates were estimated on SFM plates by inoculating plates with approximately 10^5^ spores and then harvesting after 3 and 4 days of growth. This resulted in ~1.67 × 10^9^ and 5.97 × 10^9^ spores, respectively, corresponding to 14 and 16 elapsed generations in total.

### Minimum inhibitory concentration (MIC) testing

The MIC for streptomycin of laboratory isolates was determined according to the EUCAST (European Committee of Antimicrobial Susceptibility Testing) protocol ^21^. MICs were estimated by spotting approximately 10^4^ spores on SFM plates containing 0, 2, 4, 6, 8, 12, 16, 24, 32, 48, 64, 92, 128, 192 and 256 μg ml^−1^ streptomycin sulfate (Sigma, USA). Plates were incubated at 30 °C for 4 days. The MIC was set to the lowest concentration of antibiotic yielding no visible growth. To investigate the level of streptomycin resistance in nature, we determined the MIC of a collection of 85 *Streptomyces* strains isolated from soil collected from the Himalaya in Nepal and Qinling Mountains in China ^22^. MICs were estimated as described above by spotting 1 ul of a 100-fold diluted spore stock.

### Spontaneous streptomycin resistance

Spontaneous streptomycin resistant clones were isolated from the WT strain by plating 10^9^ spores onto SFM agar containing 2, 4, 8 or 16 μg ml^−1^ streptomycin. After 2-3 days of growth, random single colonies were selected from independent plates from each streptomycin concentration and then restreaked onto a plate containing the same concentration of streptomycin as the selection plate. Spore stocks of these single colonies were collected as outlined above and stored at −20°C.

### Experimental evolution at sub-MIC streptomycin

To investigate the evolution and costs of streptomycin resistance at sub-MIC concentrations of streptomycin, we serially transferred six replicate populations for ~500 generations on plates containing 0.2 μg ml^−1^ streptomycin. This value corresponds to the minimum estimate of the Minimal Selective Concentration (MSC) for spontaneous resistant clones and is ~1/10 the MIC of the susceptible parent strain. Replicate populations, initiated from independent colonies, were grown for either 3 (14 generations) or 4 days (16 generations), after which spores were harvested as above, and then replated at a density of approximately 10^5^ spores/plate. Experimental populations were stored at – 20 °C after every transfer. After ~332 generations replicates of all six populations were in addition serially transferred to plates containing 0.4 μg ml^-1^ streptomycin, leading to a total of 12 populations. To quantify the evolution of streptomycin resistance through time we plated 10^5^ spores of all evolved populations at 50-generation intervals onto SFM supplemented with 2 μg ml^-1^ of streptomycin. Resistant colonies were scored after 6 days of growth. After ~500 generations a single random resistant colony was isolated from each 0.2 μg ml^-1^ population to be used to quantify the fitness of evolved resistant clones. This same clone was subsequently sequenced.

### Fitness assays

To assess the fitness of the spontaneous and evolved streptomycin resistant strains, we carried out head-to-head competition experiments between evolved clones and ancestral clones that were differentially marked with an apramycin-resistance cassette ^23^. Costs of resistance were quantified by competing strains in the absence of streptomycin, while the MSC of resistant clones was determined by competing strains in the presence of 0, 0.125, 0.25, 0.5 and 1.0 μg ml^-1^ streptomycin (susceptible clones at or above the MIC were fully displaced). Competition assays were initiated by mixing strains 1:1 and then plating 10^5^ total spores onto SFM at the indicated streptomycin concentration. To determine the fraction of the inoculum that was apramycin resistant or sensitive, we simultaneously plated a 10^-3^ dilution of this mix on SFM and SFM containing 50 μg ml^-1^ apramycin sulphate (Duchefa Biochemie, The Netherlands). After 4 days of growth at 30 °C the plates were harvested and the numbers of each competitor quantified following plating on SFM agar plates with or without 50 μg ml^-1^ apramycin. Control assays between WT and WT_Apr_ ancestral clones were used to correct for any fitness effects associated with the apramycin marker. Following Lenski et al (1991), relative fitness was calculated as the ratio of the Malthusian parameters of both strains: *w* = ln[x(t = 4)/x(t = 0)]/(ln[α(t = 4)/α(t = 0)]), where x is the competing streptomycin resistant strain and α is the wild type or ancestral control strain and t is the time in days of growth after inoculation. For determination of the minimal selective concentration (MSC) the selection rate constant (r) was used to define relative fitness, where instead of the ratio, we calculated the difference in the Malthusian parameters of both strains ^24^. Selection rate constant was used to control for the fact that under antibiotic exposure one or both competing clones may decline in density during the course of the assay. The MSC was estimated as the antibiotic concentration where both strains have equal selection rate constants ^6^.

### DNA extraction and sequencing

Streptomycetes to be sequenced were grown in liquid culture containing 50% YEME/50% TSBS with 5 mM MgCl_2_ and 0.5% glycine at 30 °C, 250 rpm for 2 days. After centrifugation the pellet was resuspended in TEG-buffer with 1.5 mg ml^-1^ lysozyme and after 1 hour of incubation at 30 °C the reaction was stopped by adding 0.5 volume of 2M NaCl. DNA was extracted using standard phenol/chloroform extraction, followed by DNA precipitation and washing in isopropanol and 96% ethanol. Dried DNA was resuspended in MQ water and then treated with 50 ug ml^-1^ of RNase and incubated at 37 °C for 1 hour. Following RNase treatement, the mixture was purified and cleaned as above, after which the purified DNA was washed with 70% ethanol and resuspended in MQ water. The genomes of the spontaneous and evolved clones as well as those of their ancestral strains were sequenced on the Illumina HiSeq4000 with paired-end 150 bp reads at the Leiden Genome Technology Center (LGTC). All samples were prepped with an amplification free prep (KAPA Hyper kit) after Covaris shearing of the DNA.

### Sequence analysis

All genomes were assembled to the *S. coelicolor* A3(2) genome sequence available from the NCBI database (http://www.ncbi.nlm.nih.gov/assembly/GCF_000203835.1/) using Geneious 9.1.4. The ‘Find variations/SNPs’ tool in Geneious was used to identify SNPs and indels with a minimum sequencing coverage of 10 and a variant frequency of at least 50%. Unique mutations in the spontaneous and evolved resistant strains were identified by direct comparison with the ancestral strains.

## Results

### Streptomycin resistance among natural isolates

To assess the level of streptomycin resistance among streptomycetes in nature, we tested the MICs of 85 natural *Streptomyces* strains originally isolated from the Himalaya and Qinling Mountains ^22^. In accordance with literature estimates we found resistance in a substantial fraction of these strains (46%) with low level resistance being more prevalent than high level resistance ^25^. This survey confirms that streptomycin resistance is common among streptomycetes in nature and raises questions about the benefits of streptomycin resistance at the presumably low streptomycin concentrations in the soil. Here we use the well-characterized lab strain *Streptomyces coelicolor* M145 that, with an MIC of 2 ug/ml streptomycin, has negligable resistance to streptomycin, to study the costs and benefits of streptomycin resistance.

### Spontaneous streptomycin resistance

To gain insight into fitness effects of streptomycin resistance, we isolated 16 independent clones resistant to at least 2 μg ml^-1^ streptomycin (the MIC of the susceptible WT parent strain) (Table 1). The resultant clones had MICs ranging from 4 to 192 μg/ml streptomycin (Fig. 2). Competition experiments between these resistant clones and their susceptible parent in a drug-free environment revealed that although there is significant heterogeneity in the cost of resistance (ANOVA: F_15_ = 2.92, p = 0.002), 12 of 16 resistant strains were significantly less fit than the parent, with an average cost of approximately 21% (mean ± SEM = 0.79 ± 0.018). Notably, two highly resistant clones with MICs of 196 μg ml^-1^ streptomycin appeared to have no evident costs of resistance (p > 0.05 for both clones). Across all mutants with significant costs, we found that there was no significant relationship between MIC and fitness (p > 0.05).

**Table 1.**
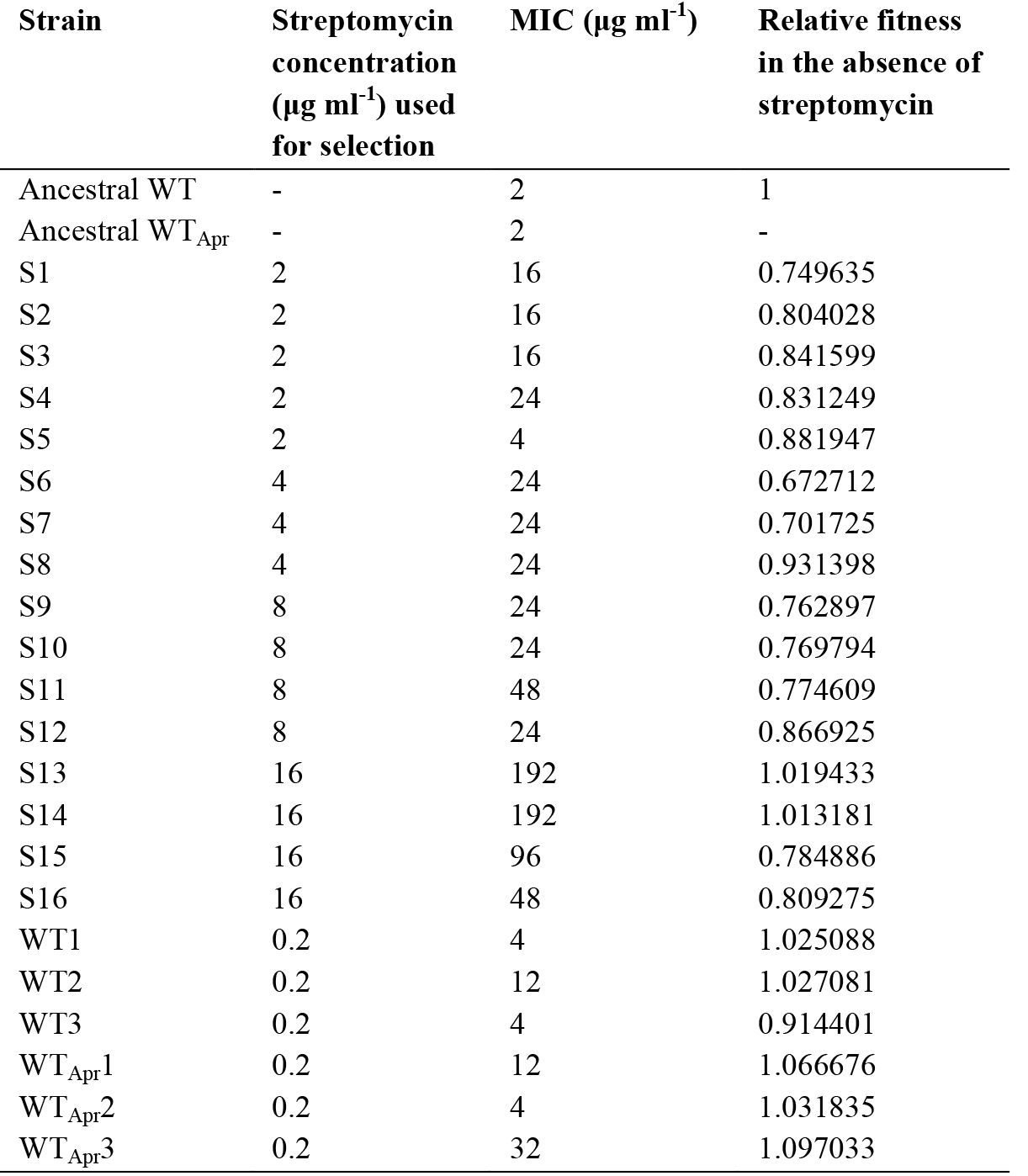
Strains used in this study

**Fig. 1.**
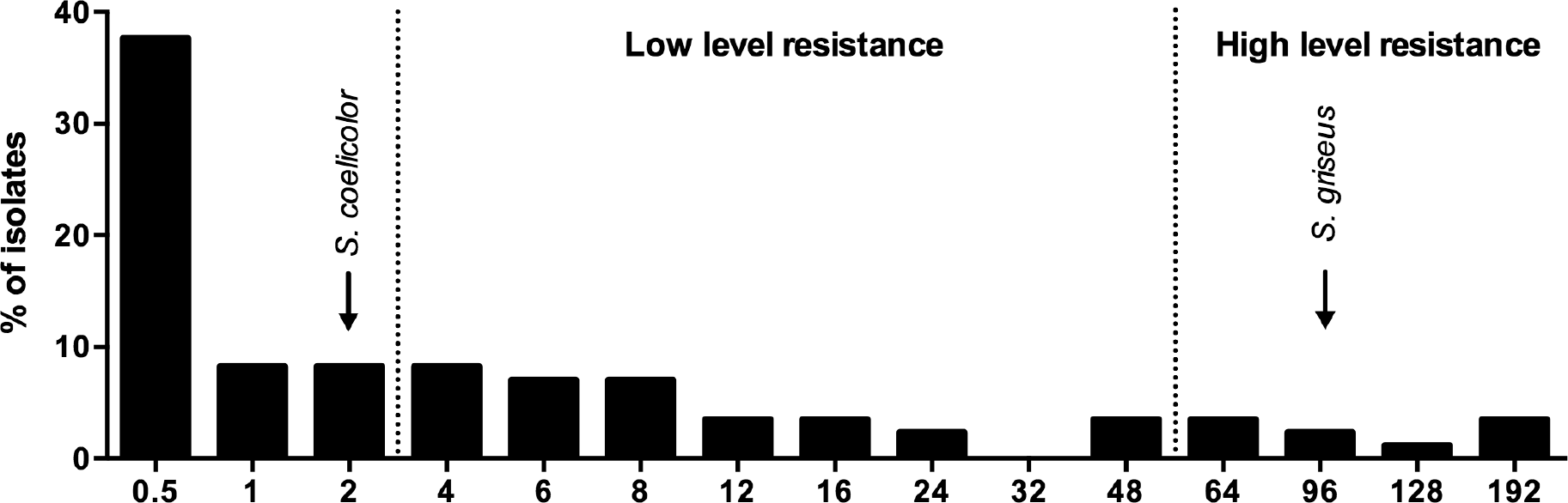
Streptomycin resistance of a collection of 85 natural *Streptomyces* isolates. MICs of *S. coelicolor* and *S. griseus* are indicated in the figure.

**Fig. 2.**
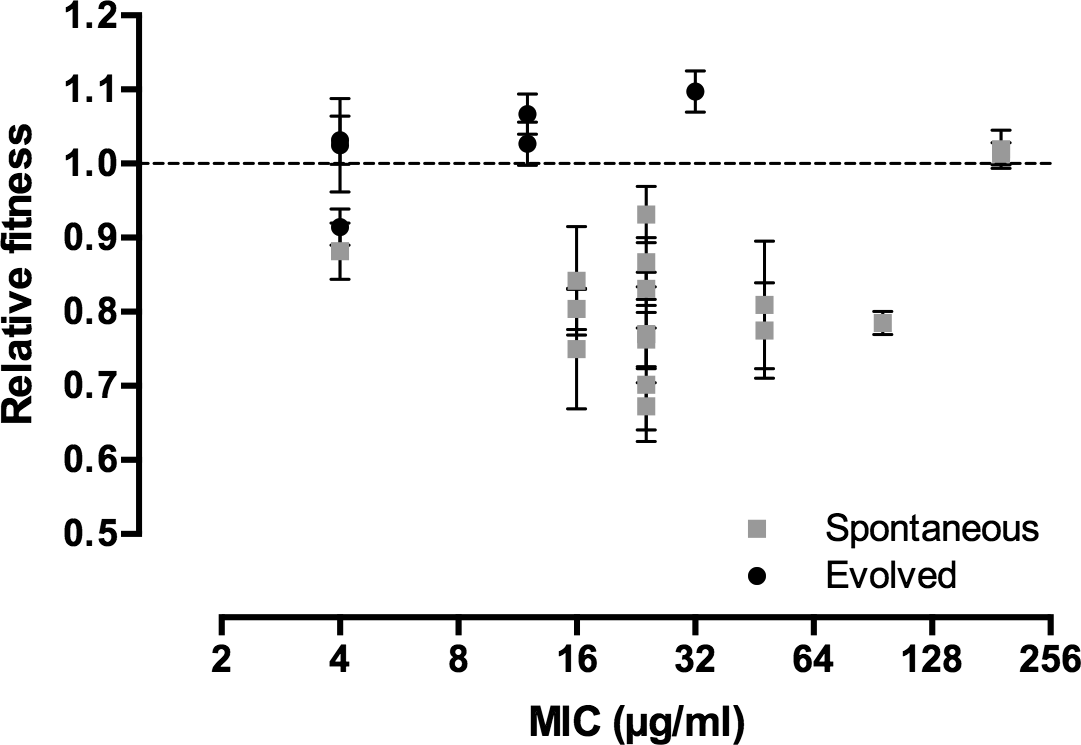
Relative fitness in the absence of streptomycin as a function of the MIC for the spontaneous and evolved streptomycin-resistant mutants. Error bars represent standard error of the mean.

To estimate the Minimal Selective Concentration (MSC) we carried out competition experiments for a subset of clones across the breadth of streptomycin MIC at increasing streptomycin concentrations and determined the MSC as the antibiotic concentration where the fitness of the susceptible and resistant strain are equal. Figure 3 shows the change in fitness as a function of streptomycin concentration for seven strains, from which we draw two conclusions. First, the fitness of each strain is strongly dependent on the drug concentration to which it is exposed during competition; as anticipated, fitness is lowest in the absence of drugs but increases sharply with small increases in the concentration of streptomycin. Second, there is variation in the MSC of different clones; the lowest MSC we measured (0.202) corresponds to ~1/10 the MIC of streptomycin against the susceptible parent strain while the highest value (0.386) corresponds to ~1/5 the MIC. These data led to the prediction that selection of *de novo* resistance should be possible at concentrations significantly less than the MIC of wild-type cells.

**Fig. 3.**
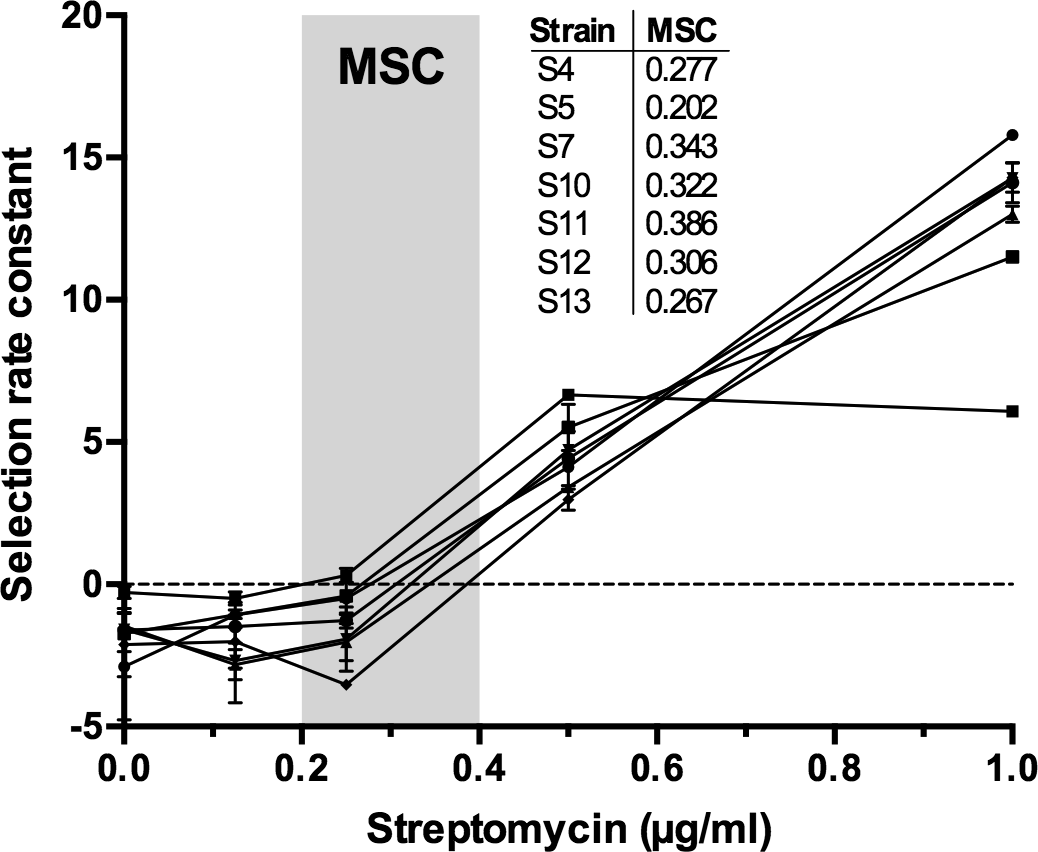
Selection rate constants as a function of the streptomycin concentration for a subset of spontaneous mutants. Error bars represent standard error of the mean.

### Evolution of resistance at sub-MIC concentrations of streptomycin

Having shown that antibiotic resistant clones gain significant fitness benefits even at low antibiotic concentrations, we next sought to determine if these same low concentrations could select for *de novo* resistance. We further aimed to quantify the spectrum of fitness costs of evolved resistant strains, as these are predicted to be lower than the costs of spontaneous resistance. We serially transferred six replicate populations on media containing 0.2 μg ml^-1^ streptomycin, which corresponds to 1/10 of the MIC of the susceptible parent strain. As shown in Fig. 4, while the frequency of resistant clones increased by at least 10-fold in three populations, with fixation of resistance in one of the populations, the remaining three populations remained static. We considered two alternative explanations for the apparent absence of resistance in these populations: either resistance mutations had not yet arisen, or alternatively, mutants were present but they were only slowly increasing due to limited benefits at the streptomycin concentrations they faced. To distinguish these possibilities we doubled the drug concentration to 0.4 μg ml^-1^ after ~ 300 generations and then continued transferring these six new populations in parallel with the original replicates. Consistent with the idea that resistant clones were present, but only slowly increasing, we observed a rapid and significant overall increase in the fraction of resistant cells in these supplemented populations as compared to those evolved at the lower concentration (paired t-test, df = 5, p = 0.028). We confirmed the evolution of *de novo* streptomycin resistance by measuring the MIC of random clones isolated from evolved populations; clones from all six populations evolved at 0.2 μg ml^-1^ streptomycin had MIC > 2 μg ml^-1^ (Figure 2).

**Fig. 4.**
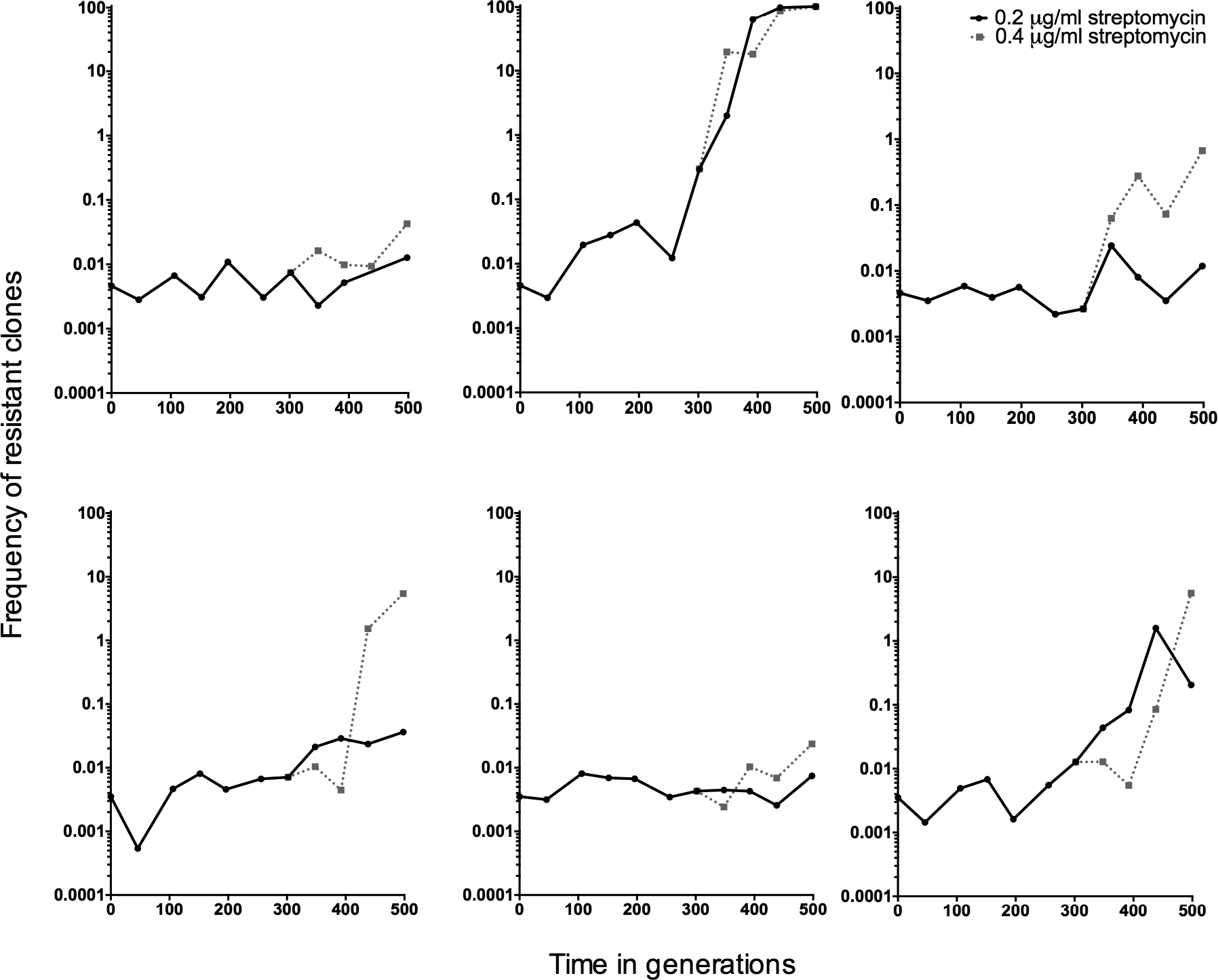
The frequency through time of strains resistant to 2 μg ml^-1^ streptomycin in populations evolved for 500 generations in the presence of 0.2 μg ml^-1^ or 0.4 μg/ml (started at ~332 generations from the 0.2 μg ml^-1^ population) streptomycin. Resistance was estimated approximately every 50 generations.

Evolution of drug resistance below the MIC is predicted to enrich for strains with reduced fitness costs of resistance. This is because resistant strains must still compete with susceptible strains that are inhibited, but not killed, by the antibiotic. To test this prediction we measured the fitness of random resistant clones in the absence of streptomycin that were isolated from the final time point of all replicated populations evolved at 0.2 μg ml^-1^ streptomycin. As shown in Figure 2, the fitness of evolved resistant clones is significantly different from the spectrum of fitness effects of spontaneous mutants (GLMM, p < 0.001). While 1 of 6 clones does have fitness costs, the fitness of the remaining five populations is either higher than or indistinguishable from 1. Overall, in contrast to the significant ~21% cost of spontaneous resistance, clones that evolved resistance at sub-MIC streptomycin had an average fitness benefit of ~3%, which did not differ significantly from 1 (Figure 2). In summary, strains of *S. coelicolor* evolving at sub-MIC streptomycin can evolve high levels of resistance while simultaneously avoiding the costs associated with this phenotype.

### Genetics of resistance

To gain insight into the mechanisms of resistance, we sequenced the genomes of ancestral and resistant strains. Across all resistant strains, we identified a total of 93 mutations: 4 synonymous substitutions, 27 non-synonymous substitutions, 3 insertions, 14 deletions (11 single bp deletions) and 45 intergenic mutations. Consistent with extensive convergence across clones, these 93 mutations mapped to only 24 genes (Table 2) and 20 intergenic regions (Table S1). On average we identified 3.1 mutations in the spontaneous mutants, with 1.6 mutations in genes and 1.4 mutations in intergenic regions. As the evolved clones were exposed to sub-MIC levels of streptomycin for 500 generations, it is not surprising that we found significantly more mutations in this set, with an average of 7.4 mutations per clone (3.7 mutations in genes and 3.7 in intergenic regions).

**Table 2.**
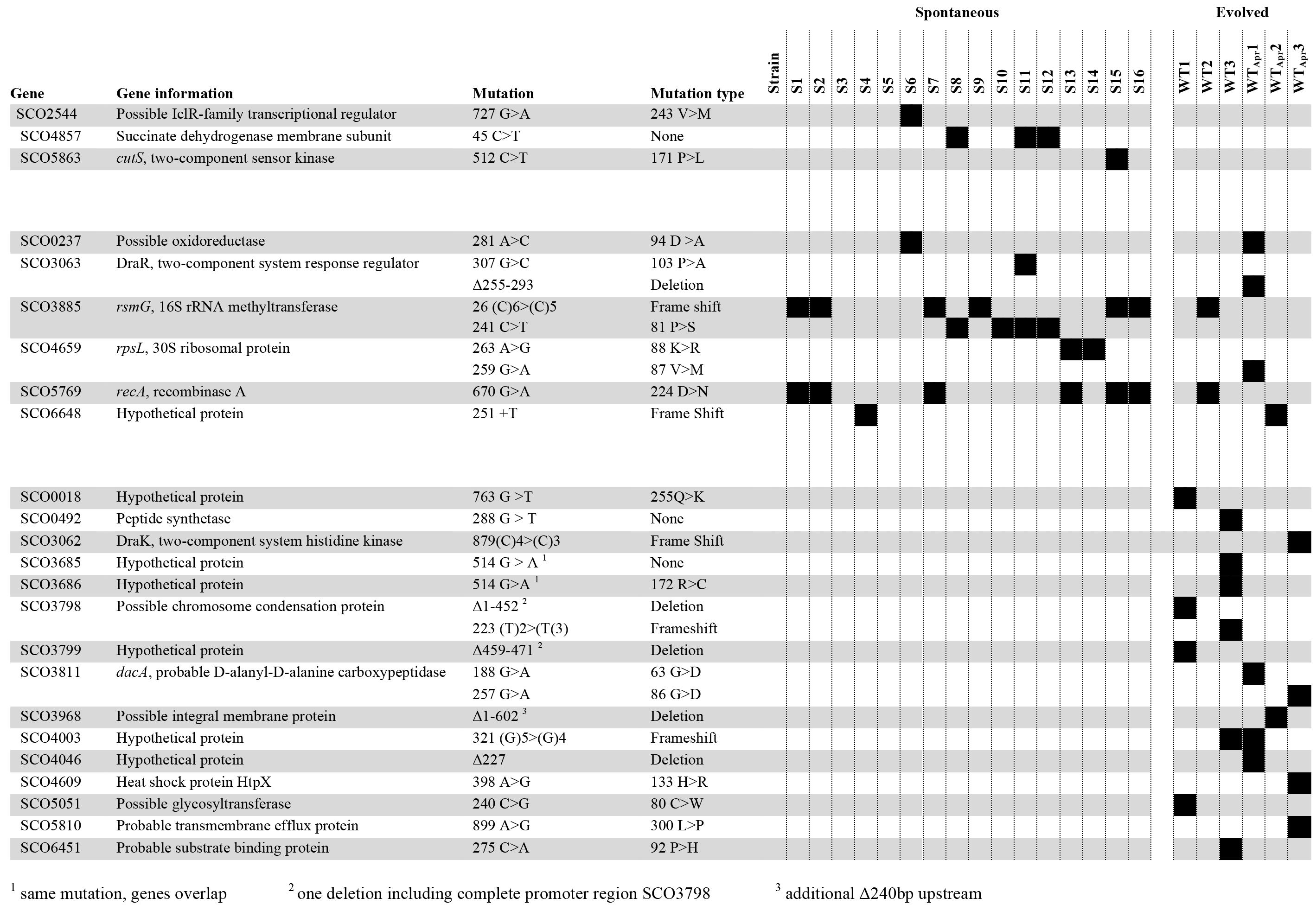
Mutations in genes in the spontaneous and evolved clones

Since the spontaneous mutants show significant fitness defects, we hypothesized that the mutations identified in this set will be costly resistance mutations, while for the evolved clones we expected to find either the same costly mutations together with others that compensate for these costs or entirely different cost-free resistance mutations. According to our results both outcomes could have occurred in our evolved lineages. Parallel mutation fixation was observed for nine genes. Six of these genes were mutated both in spontaneous and evolved mutants, strongly suggesting that they are associated with streptomycin resistance. Mutations in two of these genes, *rsmG* and *rpsL*, are known to confer low^26^ and high-level^27^ streptomycin resistance in *S. coelicolor*, respectively. Fourteen strains showed a mutation in either gene, while no strains were mutated in both genes. Eleven strains (10 spontaneous and one evolved), with MICs ranging from 12 to 96 μg ml^-1^, were found to have a mutation in *rsmG*, which encodes a rRNA methyltransferase that methylates base G527 in the 16S rRNA ^28^. Seven of these carried the same effective lesion in a homopolymeric tract of 5 cytosine residues in this gene (26 (C)6>(C)5), resulting in a frame-shift mutation that leads to an early stop codon ^26,29^, while the other four show the same non-synonymous substitution. Three clones are mutated in *rpsL*, encoding r-protein S12; one evolved clone with an MIC of 12 μg ml^-1^ and two spontaneous clones with an MIC of 192 μg ml^−1^, the latter two carrying the same 88K>R mutation that is known to cause high level resistance ^29^. Interestingly, these two spontaneous mutants (S13 and S14) are highly resistant to streptomycin, yet neither bears a cost of resistance.

While these are the only genes known to cause streptomycin resistance in streptomycetes, the fact that parallel mutations were fixed elsewhere, suggests that these mutations may be causally associated with streptomycin resistance. An interesting case can be made for the two-component system consisting of response regulator DraR (SC03063) and sensory kinase DraK (SC03062). While two strains (one spontaneous and one evolved) showed a different mutation in the gene for DraR, another strain was mutated in the gene for DraK. The DraR two-component system has been shown to be involved in the regulation of antibiotic production in *S. coelicolor* and the structural configuration of the extracellular signal domain of DraK is pH dependent, but its ligand is not known ^30,31^. Surprisingly, seven resistant strains have the same mutation in *recA*, encoding recombinase A that is involved in the homologous recombination of single stranded DNA. This mutation always cooccurs with a mutation in *rsmG* or *rpsL*; however, when comparing strains that do not have this additional mutation in *recA* we do not see a difference in MIC or fitness, implying that it may not be involved in streptomycin resistance or compensatory mechanisms. 0ther parallel mutations occuring both in spontaneous and evolved strains were located in a possible oxidoreductase and another hypothetical protein.

Two out of six evolved strains share no mutations with those arising in spontaneous resistant strains, suggesting that resistance in these strains has a different origin. Within the mutations appearing only in the evolved clones, there are three cases of parallelism. A possible chromosome condensation protein is mutated in both of the evolved strains that do not share any mutation with the spontaneous mutants, making it a likely candidate for conferring streptomycin resistance. Two evolved clones are mutated in *dacA*, which encodes a D-alanyl-D-alanine carboxypeptidase, an enzyme belonging to the group of penicillin binding proteins involved in cell-wall synthesis. Notably, we identified a mutation in the promoter region of the same gene in a third evolved clone, 101 bp upstream of the predicted translational start site. The third parallel mutation is located in a hypothetical protein. Furthermore, we identified mutations in 12 more genes that were only mutated in evolved clones, none of which were shared with the spontaneous resistant isolates.

## Discussion

Despite the appropriate emphasis on the clinical crisis in antibiotic resistance, it is also important to recognize that antibiotic resistance is a natural phenomenon that long predates the modern selective pressure of antibiotic use by man ^32^. Genes for antibiotic resistance are commonly found in nature ^33^, even in pristine environments untouched by human influence ^34,35^; however, very little is understood about the processes by which antibiotic resistance arises in these conditions. This has led to questions about the role of antibiotics in soil, where their concentrations are believed to be extremely low, as well as the role of resistance at these sub-lethal concentrations ^9,36^. Here we focus on the evolution of antibiotic resistance in the soil bacterium *S. coelicolor* in response to streptomycin, an antibiotic produced by *S. griseus*. We first show that streptomycin resistance among natural *Streptomyces* isolates is widespread, with approximately 50% of strains reaching an MIC greater than *S. coelicolor*. Next we show that costly antibiotic resistance can be offset at very low streptomycin concentrations; drug concentrations of antibiotics as low as 1/10 the MIC of susceptible strains are sufficient to provide direct fitness benefits for resistant strains. Using experimental evolution, we next find that resistant strains readily evolve during evolution at very low concentrations and furthermore that these evolved mutants are cost-free, in striking contrast to strains that evolved spontaneous resistance that carried fitness costs of more than 20%. Finally, whole genome sequencing revealed that sub-MIC evolved mutants contained a distinct spectrum of mutations from strains emerging at high concentrations.

There are several important implications of these results. First, consistent with the results of Gullberg et al (2011), our data clarify that antibiotics do not need to reach lethal concentrations to exert pronounced effects on resistance evolution. Even if susceptible cells are not obviously inhibited by sub-MIC antibiotics, their growth rates are diminished and this provides a broad range of opportunities for resistant cells to increase in frequency ^5,6,37^. This has clear relevance to the evolution of resistance in soil, where antibiotic concentrations due to endogenous production by microorganisms, both bacteria and fungi, are believed to be typically too low to inhibit competing susceptible strains ^4,8,38^. Thus, even if antibiotics are produced at sub-lethal levels, a suggestion requiring further study, they can nevertheless strongly and directly select for the emergence of resistant cells. Accordingly, emphasis on the MIC of bacteria is likely to be misguided for understanding the roles of both antibiotic production and resistance in soil; instead, emphasis should be reoriented to the MSC in order to determine boundary conditions for the emergence of resistant isolates.

Second, our results have clear implications for the persistence of resistant strains. Resistant bacteria that were isolated following exposure to lethal streptomycin concentrations were burdened with significant fitness costs, an outcome widely observed across species ^39^. One possibility is that these effects are caused by resistance mutations, e.g. in *rpsL* or *rsmG*, that lead to hyper-accurate protein translation and therefore slower growth ^40,41^. Alternatively, and specific to Streptomyces,streptomycin resistance can in some cases lead to hyper-production of antibiotics ^27,29,42^, although we did not observe any increased susceptibility of our ancestral strain of *S. coelicolor* to any of the evolved strains. In contrast to bacteria that were selected at high streptomycin concentrations ^39,40^, *S. coelicolor* strains that evolved resistance at sub-MIC doses were cost-free. From an environmental standpoint, this suggests that resistance evolving at sub-MIC antibiotics in soil will persist in the face of competition with susceptible cells, while cells that bear the significant fitness costs of spontaneous resistance would be predicted to decline ^8,17^. Consistent with this, and as observed in more detail here, streptomycin resistance is commonly found in nature, with low-level resistance being more prevalent than high-level resistance ^25,43^. Although there are many potential reasons for this, including high densities of *S. griseus* that are naturally resistant to their own antibiotic^18^, resistance in other species may arise because of the direct benefits resistance provides. From a clinical standpoint, cost-free mutations emerging at sub-MIC antibiotic concentrations are problematic because this could serve to reduce the reversibility of resistance, a potential that relies on durable fitness costs in resistant isolates^17^. Certainly, infectious bacteria face a range of antibiotic doses during treatment ^5,44^; if this influences the types of resistance mutations that arise and fix, and in particular their costs, it will be necessary to take this into consideration during the development of treatment protocols.

Third, our results suggest that resistance mutations selected at sub-MIC concentrations are distinct from those arising above the MIC. While mutations in genes *rsmG* and *rpsL*, known to be associated with streptomycin resistance, were identified in 12 out of 16 spontaneous and 2 out of 6 evolved clones, the resistance mechanisms in the other clones remain to be elucidated. Many of the mutations/mutated genes occur in parallel, suggesting that they are directly involved in streptomycin resistance or potentially that these mutations influence the costs of resistance. For example, the DraR-K two-component system is mutated in several lineages. Various two-component systems have been implicated in the control of antibiotic production^45^, but as far as we are aware none have been specifically tied to resistance in the absence of the related biosynthetic gene cluster. Further research into the DraR-K response regulon is required to shed light on this important phenomenon. Another intriguing parallel mutation is located in *recA* and was found in seven sequenced strains. As a disruption of *recA* in *S. coelicolor* increases genetic instability ^46^, it is possible that this mutation increases the likelihood for subsequent resistance evolution. Despite these cases of parallelism, many evolved lineages carry unique mutations in hypothetical genes or intergenic regions. Moreover, there is little overlap between mutations found in sub-MIC evolved lineages and those selected for spontaneous resistance at higher drug concentrations. This indicates that many routes and mechanisms towards drug resistance are unknown. Also, it may indicate that studying antibiotic resistance at lethal doses provides only part of the spectrum of resistance mutations. At present, the role these mutations play in resistance is unknown; however, these are strong candidate for testing in future work. In addition, these mutations clarify the value of using experimental evolution at sub-MIC drug concentrations to elucidate novel modes of resistance. Finally, we note that in 2 of 16 spontaneously resistant lineages we failed to identify any mutations at all. Although our coverage was high in these clones, the *Streptomyces* chromosome is very GC rich (>70% G+C content), making assembly challenging and rendering certain regions difficult to sequence. Additionally, short-read sequencing may fail to capture duplications that could be highly relevant for resistance evolution ^47^. Longer-read sequencing platforms should hopefully address these problems in this system in the future.

While antibiotics have been traditionally considered as inter-bacterial weapons ^38^, their role has been reexamined in the last few decades in light of results showing that cells respond to sub-MIC antibiotics with broad and diverse changes to gene expression and cellular physiology ^4^. By this new view, antibiotics are not weapons but instead are reinterpreted as signals, while resistance is understood to modify signal strength ^9,36^. We recently cast doubt on this reinterpretation in studies showing that social interactions among competing Streptomycetes had a dramatic influence on antibiotic production ^48^, a result consistent with their likely role as inter-microbial weapons. The present work supports this view. In short, irrespective of any other effects sub-MIC antibiotics have on cells, these low concentrations are sufficient to both inhibit competing susceptible cells and to provide sufficient natural selection to enrich for resistance.

Several questions nevertheless remain from this study. First, we lack a clear understanding of the effective concentrations of streptomycin in soil. While concentrations are often claimed to be low, little direct evidence supports this possibility, and local concentrations may in fact be high. Moreover, it remains unclear how antibiotic concentrations in soil are influenced by the physico-chemical properties of soils together with the role of other inter-microbial dynamics that influence antibiotic production. It therefore remains a key goal to extend this work to more natural microcosms that include structured soil, as well as including competition with the natural streptomycin producer *S. griseus*. Second, it remains unclear why *de novo* antibiotic resistance at sub-MIC streptomycin selects for cost-free mutations. 0ur genome sequencing has identified several putatively causal mutations for resistance in two well-studied genes; moreover, it has suggested candidate genes that could either compensate for costs of resistance, or alternatively could represent entirely new suites of resistance mechanisms that are intrinsically cost-free. This needs to be followed with more mechanistic studies to determine the precise functional role of these mutations. Finally, it will be important to extend our analyses to the evolution of resistance in natural environments influenced by anthropogenic antibiotic pollution ^7,8^. Natural reservoirs for resistance can transfer genes for resistance to clinically relevant pathogens ^49^; if these mechanisms are enriched for low-cost resistance mutations, then this has profound potential consequences for the distribution and persistence of resistance types among infectious bacteria.

## Acknowledgements

Financial support was provided by a grant from the Dutch National Science Foundation (NWO) to D.E.R. and by a grant from the China Scholarship Council (CSC) to Z.Z. Additional support was provided by the UK Biotechnology and Biological Sciences Research Council [BB/J006009/1] to D.E.R. and Ian S. Roberts (University of Manchester).

---

**Table S1.**
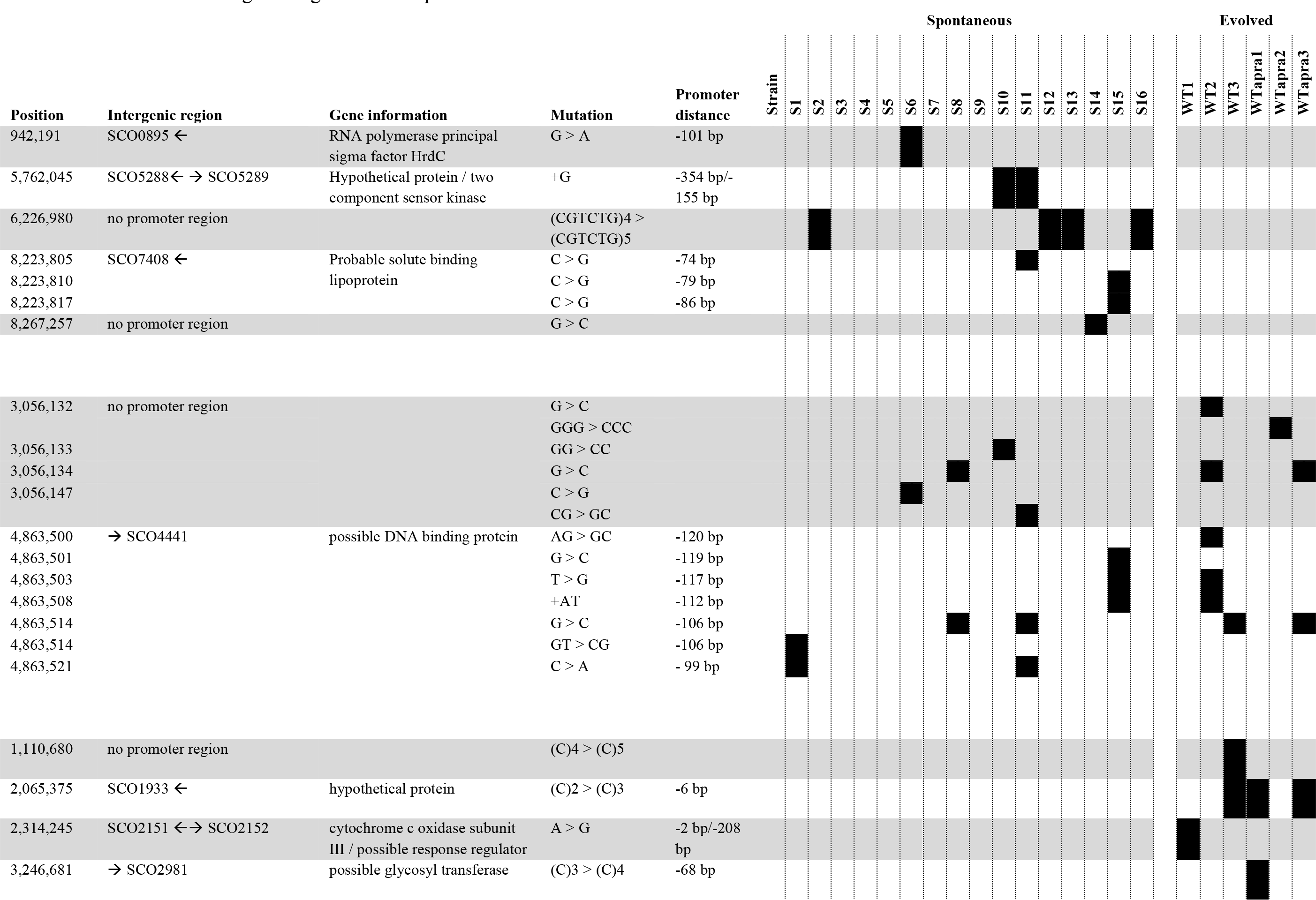
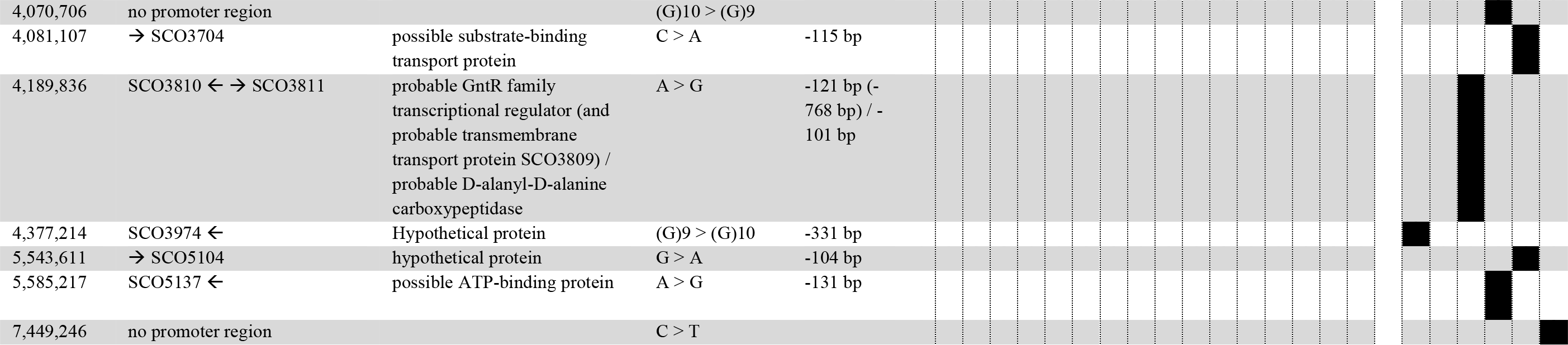
Mutations in intergenic regions in the spontaneous and evolved clones

